# Soilless system design impacts the diversity and composition of microbiota

**DOI:** 10.64898/2026.02.19.706784

**Authors:** Auja Bywater, Aline Novaski Seffrin, Jordan E Bisanz, Francesco Di Gioia, Jasna Kovac

## Abstract

Controlled environment agriculture (CEA), including soilless farming systems, is rapidly expanding as a strategy to improve food security and resource efficiency. However, limited information is available on how different soilless farming system designs influence microbial populations relevant to plant health and food safety. This study investigated the effects of soilless growing systems and growing season on aerobic plate counts (APC) and bacterial community composition in nutrient solution and on bok choy (*Brassica rapa* subsp. *chinensis*) leaves. Five soilless systems, deep water culture (DWC), Kratky (KR), nutrient film technique (NFT), ebb and flow (EF), and drip irrigation (DI), were evaluated across fall and spring growing seasons. Soilless system type significantly influenced APC in nutrient solution, with the DI system consistently exhibiting the highest counts across both seasons. Increased nutrient solution pH was negatively associated with APC, whereas temperature did not significantly affect bacterial concentrations. In contrast, APC on bok choy leaves were not significantly influenced by system type, season, pH, or temperature. Bacterial community composition in nutrient solution was strongly shaped by season, soilless system type, sampling day, and temperature, as determined by 16S rRNA V4 amplicon sequencing. Microbial diversity varied primarily by system type, with limited influence of pH or temperature. Core microbiota analysis identified a small subset of taxa that persisted across systems and seasons, with *Acidovorax* detected in all samples. We found that soilless system design and seasonal conditions strongly influence microbial load and community structure in nutrient solution, providing a foundation for developing system-specific microbial management strategies.

**Importance:** Understanding factors that shape microbial community composition in soilless farming systems is critical for optimizing plant health, system productivity, and food safety. Microbial communities influence nutrient cycling, biofilm formation, and pathogen survival, all of which affect the ecological stability and performance of these systems. By identifying how system design, seasonal variation, and environmental conditions influence shifts in microbial populations, targeted strategies can be developed to promote beneficial microorganisms and mitigate risks associated with pathogens. This knowledge contributes to advancing safe and sustainable soilless farming practices that can meet the growing demand for fresh produce grown in controlled environments.

## Introduction

Controlled environment agriculture (CEA), integrating the use of soilless farming techniques that allow for growing fresh produce without soil, is a rapidly growing agricultural sector. Soilless farming allows for growing food in areas otherwise unsuited for farming, including cities, deserts, and regions with poor soil or limited freshwater resources, which can increase food security globally (1, 2). Several alternative soilless production system designs exist, each differing in substrate utilization and nutrient solution delivery strategies. A few examples include the deep water culture (DWC), Kratky (KR), nutrient film technique (NFT), ebb and flow (EF), and drip irrigation (DI) systems (3, 4). Both the DWC and KR are static soilless systems, with plant roots suspended in a reservoir of nutrient solution, with DWC incorporating an air stone to enhance oxygenation (5, 6). The NFT system utilizes shallow channels with a small pump to circulate the nutrient solution continuously over the plant roots (7). In contrast, EF and DI systems incorporate a solid growing substrate, often a peat-based mix, to support root structure and retain moisture (8, 9). EF operates by periodically flooding and draining the root zone, while DI delivers nutrient solution at set intervals through drippers. While these systems operate differently, they all utilize substrate and/or water, both of which have been identified as potential sources of bacterial introduction (10).

Many bacteria found in soilless farming systems exhibit traits that may facilitate their persistence in water-based environments, particularly motility and biofilm formation (11, 12, 13). Motility allows bacteria to actively seek nutrients, colonize surfaces, and interact with plant roots, especially in flowing or aerated systems (14). Plant pathogens, such as *Phytium* spp., and human pathogens, such as *E. coli, Salmonella,* and *Listeria,* are examples of motile bacteria that can cause plant disease in these systems or human disease after consumption of contaminated produce, respectively (15). Biofilm formation enables microbial communities to adhere to surfaces and resist environmental stressors, including sanitizers (16, 17). These traits are advantageous for bacteria in soilless farming systems, where continuous nutrient solution flow and an absence of soil create selective pressures favoring bacteria capable of surface attachment and active movement. Biofilm forming bacteria isolated from an NFT system facilitated the attachment and colonization of *Salmonella* on PVC surfaces, further highlighting microbial food safety risks in soilless farming systems (18).

Each type of soilless growing system influences the interaction between the roots, substrate, nutrient solution, plants, and microbial populations (15). These unique setups can influence factors like oxygen level, pH, temperature, and nutrient availability, all of which are important growth and survival factors for bacterial populations. Different systems also create unique ecological conditions that shape microbial communities in ways not yet fully understood (19, 20). The microbiota in soilless farming systems can influence critical functions like nutrient cycling, biofilm formation, and suppression or proliferation of various bacteria (20, 21). These microbial processes have direct implications for plant health, yield, and food safety, and could be utilized to optimize crop management (19). The primary objective of this study was therefore to investigate the influence of alternative soilless growing system types and the associated environmental factors on microbial populations in soilless growing systems to understand how these factors shape microbial diversity and composition over time.

## Results and Discussion

### The Drip Irrigation System Had the Highest Aerobic Plate Counts Across Seasons

The average aerobic mesophilic bacteria plate counts (APC) in the nutrient solution were 4.7 ± 0.76 log_10_ and 4.4 ± 0.84 log_10_ CFU/mL in fall and spring, respectively (*p =* 0.001; Fig. 1). Hydroponic system type significantly influenced APC in both the spring and fall (*p* < 1.824e-08; Fig. 1). Furthermore, there was a significant interaction between the system and season (*p* = 0.001; Fig. 1), indicating that the effect of the soilless system type on APC varied by season. In the fall, APC concentrations ranged from 2.39 log_10_ CFU/mL (EF) to 7.47 log_10_ CFU/mL (DI). In the spring, counts ranged from 2.6 log_10_ CFU/mL (EF) to 6.43 log_10_ CFU/mL (DI). The DI system exhibited significantly higher average APC than the other systems in both seasons (*p* ≤ 0.012; fall: 5.9 ± 1.01 log_10_ CFU/mL, spring: 5.44 ± 0.6 log_10_ CFU/mL). Within DI and KR systems, APC did not differ significantly between fall and spring (*p >* 0.08). In contrast, APC was significantly higher in fall than in spring in the DWC (*p =* 0.008) and NFT (*p =* 0.001), whereas the EF system exhibited higher APC in spring compared to fall (*p =* 0.036). While the system-season effect on APC in nutrient solution was significant, it did not have a significant system impact on APC on bok choy leaves in either season (Fig. S1).

**FIG 1.**
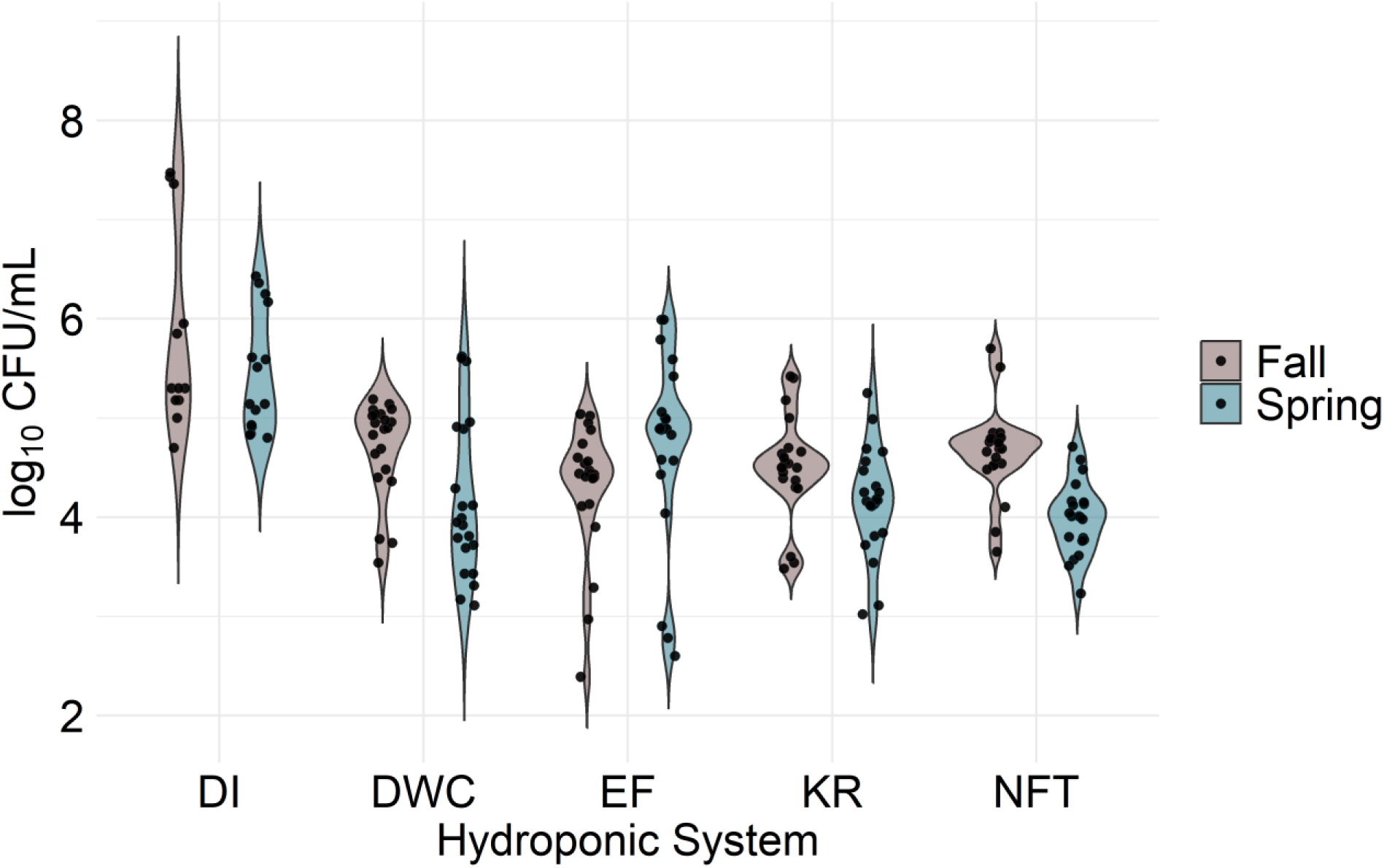
Mesophilic aerobic bacteria concentrations, as measured by Aerobic Plate Count (APC) analysis of nutrient solution sampled from different soilless growing systems in fall (n = 97) and spring (n = 98). DI, drip irrigation; DWC, deep water culture, EF, ebb and flow, KR, Kratky, NFT, nutrient film technique.

The consistently elevated APC in the DI system nutrient solution may be partially attributed to substrate composition. Both DI and EF systems contained a peat-based substrate derived from the surface organic soil layer composed of partially decomposed plant biomass, which may introduce bacteria and represent a potential source of baseline microbial input. Peat-vermiculite application in soil-based crop production has been shown to increase carbon availability and support higher bacterial populations (22). However, because only the DI system exhibited elevated APC across both seasons, substrate composition alone does not fully explain the observed differences. Instead, differences in nutrient solution delivery and retention, such as the localized moisture at the root zone in DI compared to the flooding and draining in EF, likely contributed to the observed patterns. The EF system periodically flooded the pots, with nutrient solution being absorbed primarily from the lower substrate region, potentially limiting exposure of the upper substrate to irrigation water. In contrast, the DI system applied nutrient solution at the top of the pot, allowing complete percolation through the substrate profile and greater potential contact with all substrate-associated microorganisms. Furthermore, the higher volume of water circulated through the EF system may have caused a dilution effect, lowering the concentration of bacteria per unit of volume and resulting in lower APC concentrations determined for EF compared to the DI system.

While the DI system consistently maintained the highest APC across both seasons, temporal trends varied among systems (Fig. 2). A significant interaction between system and sampling day was observed in both seasons (*p =* 5.94 x 10^9^), indicating microbial levels changed over time and these changes differed by the system. In the fall, APC values increased significantly in the NFT (*p =* 0.0171) and DWC (*p =* 0.0002) systems, with concentrations rising from 3.86 to 4.72 in NFT and from 3.69 to 4.98 in DWC. In contrast, the APC concentrations in DI (*p =* 0.0295) systems significantly decreased from 7.42 to 5.26. In the spring, the APC concentrations in samples collected from the EF system significantly increased over time (*p =* 0.0008), from 2.76 to 5.34, while the APC concentrations significantly decreased in the DI (*p =* 0.0001) system from 6.35 to 4.89. Temporal changes were not significant in the other tested systems and seasons. The decrease in APC over time in the DI system may seem contradictory, given that it consistently exhibited the highest overall counts. However, APC concentrations on day 0 were high in both seasons, potentially reflecting bacteria present in the peat-based substrate that were released into nutrient solution during initial submersion. This likely resulted in a statistically significant decline in APC after day 0.

**FIG 2.**
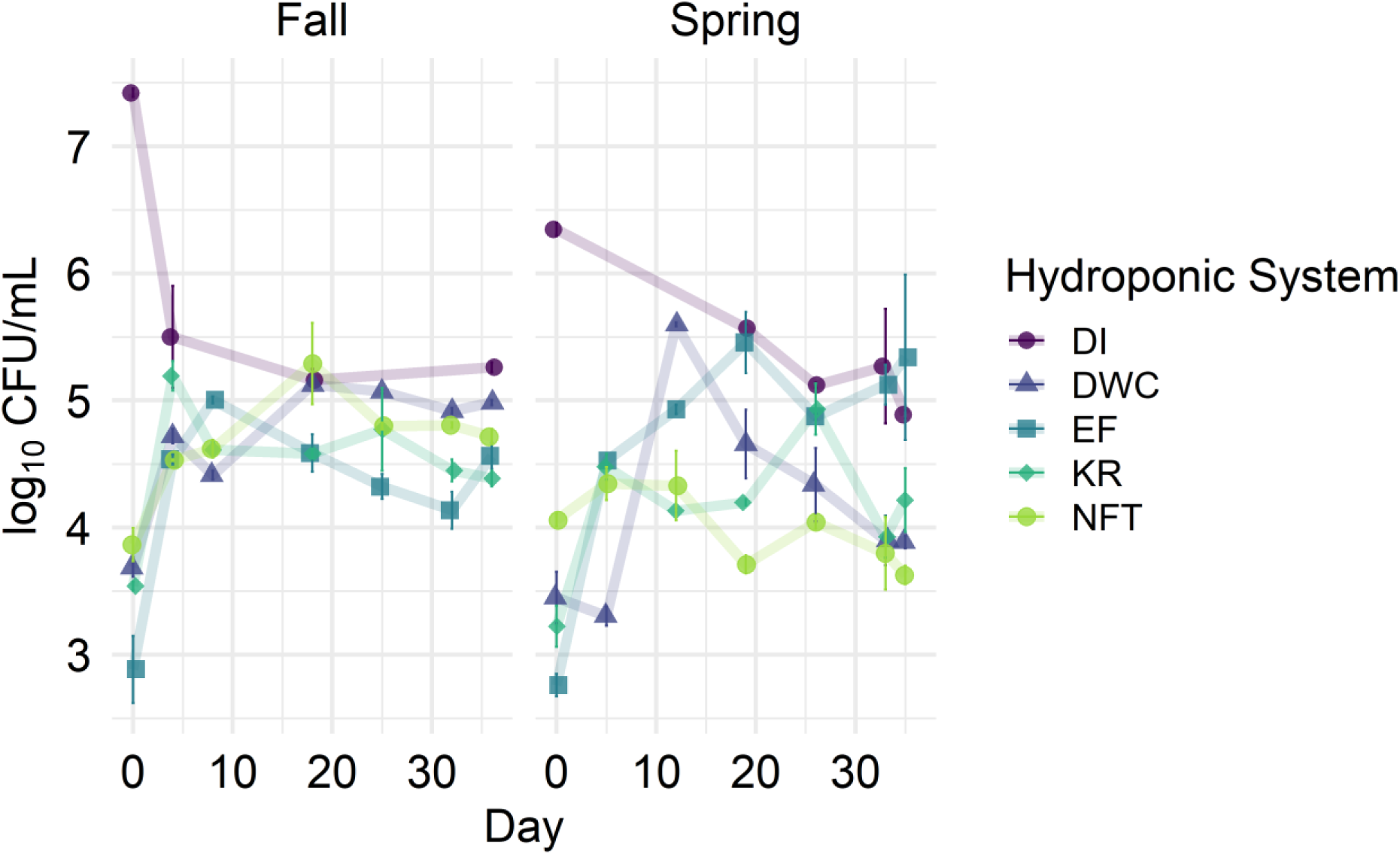
Temporal trends in aerobic mesophilic bacteria concentration, as determined by Aerobic Plate Count (APC) method, across soilless systems and between seasons. The lines connect mean APC values for each system throughout each season to indicate changes in microbial load throughout the growing cycle. DI, drip irrigation; DWC, deep water culture, EF, ebb and flow, KR, Kratky, NFT, nutrient film technique.

### PH Influenced APC Counts in the Nutrient Solution

There were several weeks when the DI system did not have enough nutrient solution run off to collect a sample for temperature and pH measurements. Due to the missing data, the DI system was removed from pH and temperature analysis. Furthermore, samples collected on day 0 were excluded from the analysis because the initial nutrient solution temperature used to fill the systems was elevated relative to subsequent time points, confounding the results by correlating higher microbial loads with lower temperatures towards the end of the growing cycle.

Nutrient solution temperature differed significantly among soilless systems (*p =* 3.55 × 10⁻⁶) and between seasons (*p <* 2.2 × 10⁻¹⁶), with no significant system × season interaction (p = 0.0618), indicating that seasonal temperature changes were consistent across systems (Table. S1). Overall, nutrient solutions were warmer in spring (22.08 ± 2.65 °C) than in fall (20.4 ± 2.3 °C). Nutrient solution pH also varied significantly among systems (*p =* 3.93 × 10⁻¹⁰) and between seasons (*p =* 3.05 × 10⁻⁷) and exhibited a strong system × season interaction (*p =* 4.17 × 10⁻⁸) (Table. S1). Across systems, mean pH was higher in spring than in fall (6.3 ± 0.7 vs. 5.9 ± 0.3) (Fig. 3).

**FIG 3.**
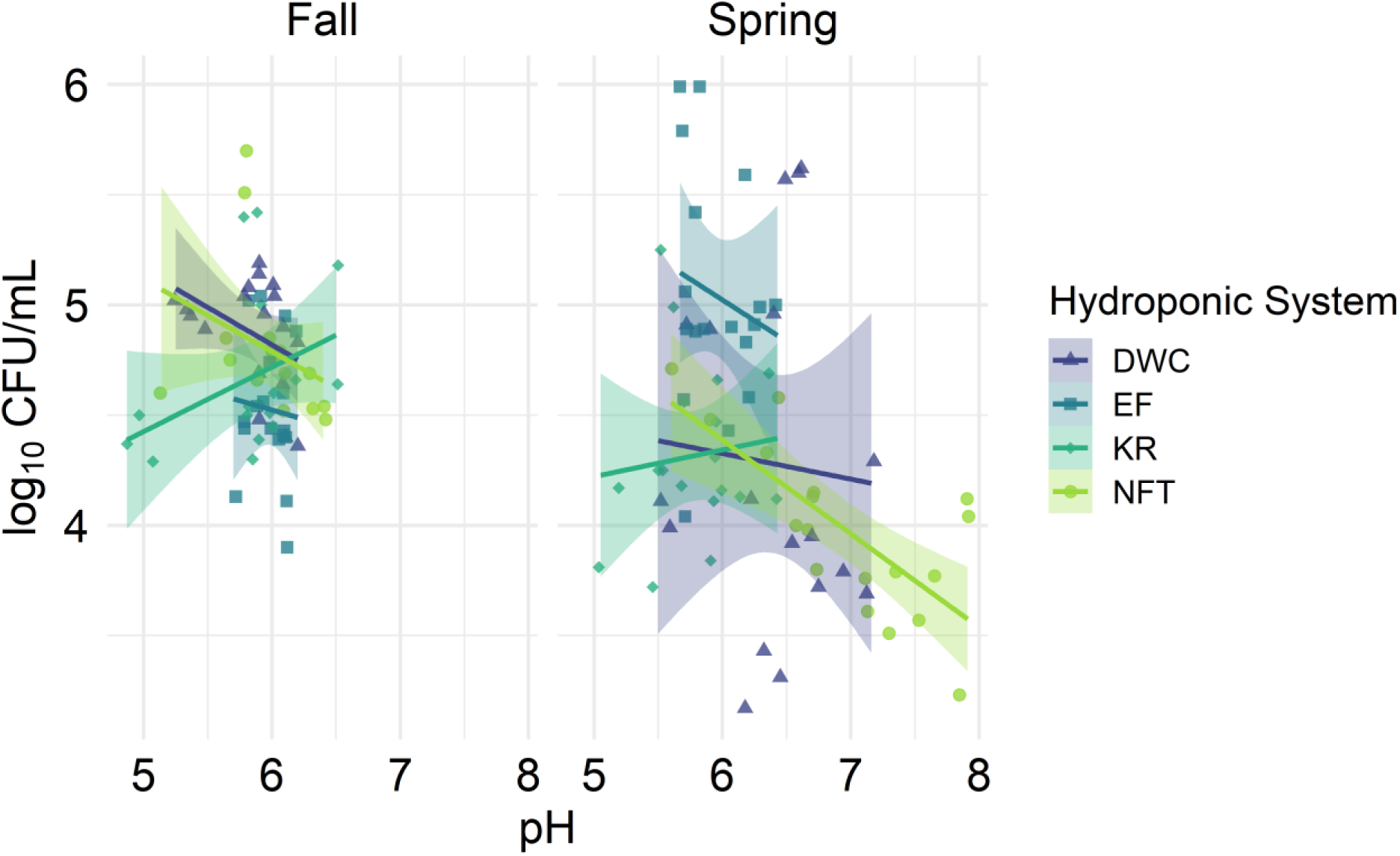
Linear relationships between pH and log_10_-transformed Aerobic Plate Counts (APC) in different soilless systems, faceted by season. DI, drip irrigation; DWC, deep water culture, EF, ebb and flow, KR, Kratky, NFT, nutrient film technique.

Temperature did not have a significant effect on APC concentration in either season (*p* = 0.47). In contrast, nutrient solution pH was significantly negatively associated with APC across systems and seasons (*p =* 3.45 × 10⁻⁵), with system-specific slope estimates indicating a consistent decrease of approximately 0.36 log_10_ CFU per unit increase in pH across all soilless systems (95% CI: –0.54 to –0.18). This finding may reflect microbial acid production associated with higher bacterial loads. Inclusion of pH in the model reduced the effect of season, indicating that some of the observed seasonal differences in APC may be partially explained by variation in pH. Neither the pH × season (*p =* 0.26) nor the system × season × pH interaction (*p =* 0.91) was significant, indicating that the relationship between pH and APC was consistent across seasons and soilless systems.

The relationship between pH and day (*p =* 0.46), as well as the day × system interaction was statistically insignificant (*p =* 0.10), indicating that pH did not exhibit systematic temporal effects within any soilless system. Although few studies have directly examined pH effects on APC in soilless systems, previous work reported that increasing nutrient solution pH from 5.0 to 6.5 increased populations of *Clavibacter michiganensis* subsp. *michiganensis* (23). In contrast, our study observed higher APC at lower pH, potentially reflecting differences in experimental context. Our study was conducted in fully operational soilless growing systems with plants, whereas Huang and Tu evaluated bacterial survival in nutrient solution alone. The association between lower pH and higher APC may reflect increased organic acid production resulting from greater microbial metabolic activity (24). There may also be a confounding effect from pH that was not captured in our data, like the impact of pH on nutrient availability in the nutrient solution. One study found that increasing or decreasing an environmental pH by as little as one unit can influence the metabolic activity of microbial communities by up to 50% (25). Our study did not assess nutrient solution compositional changes, but future studies could include this to further understand potential confounders. Soilless system type also contributed to pH variation. ANOVA followed by Tukey’s test showed that the NFT system had significantly higher pH compared to DWC, EF, and KR systems (*p <* 0.01), and the KR system had significantly lower pH than DWC (*p =* 0.03). Although pH was negatively associated with APC within systems, differences in mean APC among systems did not fully correspond to system-level pH rankings. This indicates that while APC and pH have a relationship, other factors not measured in this study likely contributed to the observed system-specific APC differences.

The APC counts from the bok choy leaves in our study were not significantly influenced by the system, season, pH, or temperature, likely because they were not in direct contact with the nutrient solution. Other research has reported higher bacterial loads on external leaves in contact with irrigation water in hydroponic systems in Puerto Rico (26). In our study, the outer leaves had minimal or no contact with the water, likely leading to a lack of significant influence.

### Soilless System and Seasonal Conditions Drive Bacterial Variation

Baby bok choy samples were excluded from further analysis due to high prevalence of chloroplast and mitochondrial sequences resulting from cross-reactivity with common 16S V4 primers.

Season had the strongest influence on bacterial community composition in the nutrient solution (R² = 0.053, *p =* 0.001), with system type also contributing significantly (R² = 0.254, *p =* 0.001). Their interaction was likewise significant (R² = 0.105, *p =* 0.001), indicating that the effect of system type differed between seasons (Fig. 4). Sampling day also had a significant effect on bacterial community composition (R² = 0.05, *p =* 0.001), indicating that temporal changes within a season contribute to variation in the microbiota. Unlike APC concentrations, temperature significantly shaped community composition (R² = 0.045, *p =* 0.001), whereas pH did not (*p =* 0.125).

**FIG 4.**
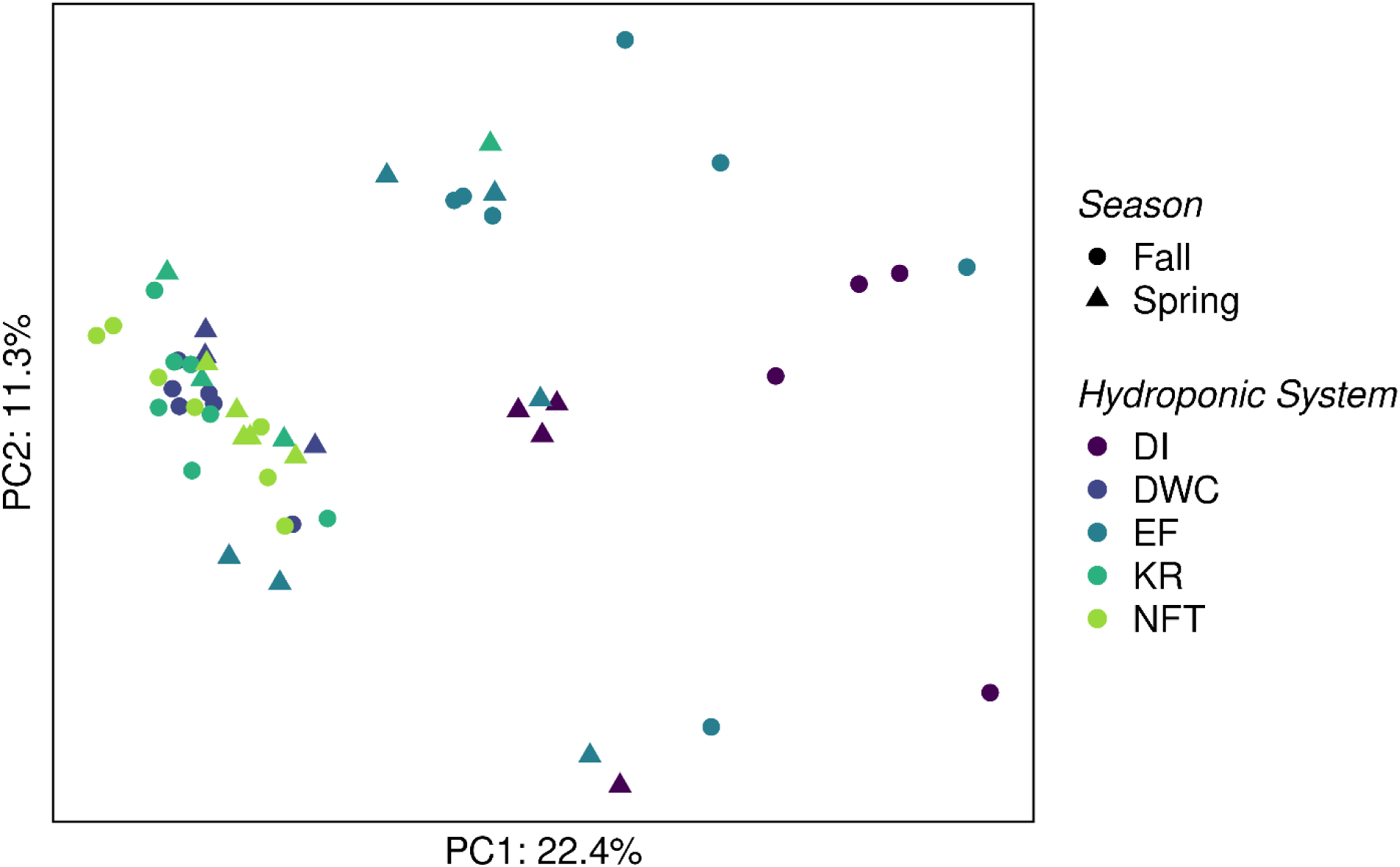
Differences in microbiota composition among systems and between seasons. PC1, principal component 1; PC2, principal component 2. DI, drip irrigation; DWC, deep water culture, EF, ebb and flow, KR, Kratky, NFT, nutrient film technique.

Average nutrient solution conditions observed in our experiment (22.05 ± 2.04 °C, pH 6.3 ± 0.63) support growth of most environmental and soilless farming associated bacteria (19, 27). However, there were still significant influences from season, temperature, system, and sampling day. The significant impact of system type on community composition was also found in another study that collected nutrient solution samples from four system types (NFT, DWC, EF, and vertical drip) across 10 commercial facilities in Ohio (20). This suggests that the system type plays an important role in community composition in larger commercial scale systems than those in our study and in different geographical regions.

In the fall, bacterial community composition differed significantly between all systems, with the exception of DWC vs. NFT (p = 0.26). In the spring, fewer differences were observed: DI differed significantly from all other systems (p < 0.03), and EF differed from NFT (p = 0.01), indicating that seasonal variation reduced the magnitude of compositional differences among systems.

To assess temporal trends in alpha diversity across different soilless systems during fall and spring seasons, linear regression models were fitted to Shannon diversity index data (Fig. 5). Due to the temperature of the starting nutrient solution being elevated compared to the rest of the samples collected throughout the experiment, we excluded data from day 0 to mitigate confounding results. When data for both seasons were combined, the Shannon diversity was not significantly affected by time, pH, nor temperature. System type was the strongest predictor, explaining most of the variation in alpha diversity across samples. In contrast to APC concentrations, pH had an insignificant effect on Shannon diversity (p = 0.974), suggesting that while pH influenced the load of mesophilic aerobic bacterial, it did not influence overall community diversity. Another study investigating microbial ecology in four different commercial soilless systems (NFT, DWC, EF, and vertical drip) growing lettuce found that temperature and pH both influenced bacterial variation in lettuce leaves, roots, growth media, and nutrient solution samples (20). The difference in pH finding could be due to the inclusion of leaf, root, and growth media samples as well as difference in the size of commercial facilities compared to our small-scale experimental set-up.

**FIG 5.**
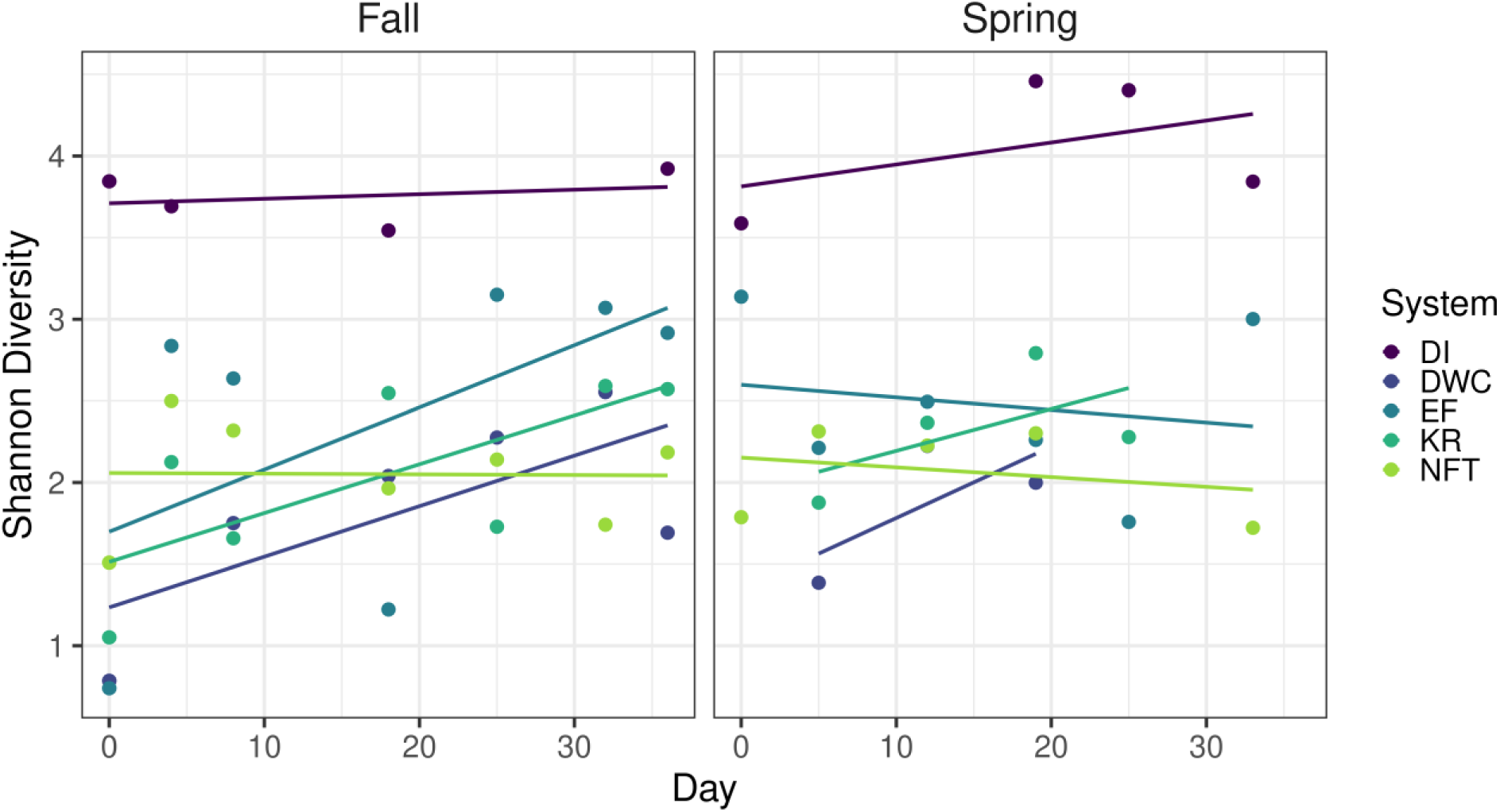
Shannon diversity index temporal trends in soilless systems in fall and spring seasons. DI, drip irrigation; DWC, deep water culture, EF, ebb and flow, KR, Kratky, NFT, nutrient film technique.

### Core and Hub Taxa were Identified Across Systems and Season

Core microbiotas were identified to determine which taxa were consistently present in the soilless systems and seasons. Core taxa were defined as ASVs detected in all tested samples within a system, and core richness varied widely among systems. Negative extraction and PCR controls were included and monitored for potential contamination. The DI system exhibited the highest core richness (292 ASVs), while the DWC, EF, and KR systems each contained 15 core ASVs, and the NFT system contained 11. Despite this variability, one taxon was shared across systems and seasons (Fig. 6),suggesting it may be a well-adapted member of hydroponic environments. *Acidovorax* was the only taxon detected in every tested sample and had mean relative abundances ranging from 0.14% (DI, fall) to 7.3% (NFT, fall). The consistent and widespread presence of *Acidovorax* underscores its persistence and potential ecological compatibility with soilless systems nutrient solutions, pH, temperature, and other microbiota in these systems. *Acidovorax* are Gram-negative and motile bacteria, which supports their dispersal, surface colonization, and resilience in water-based, low organic carbon environments (28). While nutrients in nutrient solution are abundant, the organic carbon is typically very limited, which may result in enhanced chemotaxis of heterotrophic bacteria towards root exudates and potential biofilm formation on root surfaces (29). Some *Acidovorax* strains are capable of nitrate-dependent Fe(II) oxidation coupled with organic carbon utilization, a mixotrophic lifestyle that may facilitate their persistence in carbon-limited soilless farming environments (30, 31). These characteristics likely support their competitive success in recirculating systems where movement through the nutrient solution, rather than soil structure, governs microbial interactions. Although several *Acidovorax* species (e.g., *A. avenae*, *A. citrulli*, *A. cattleyae*) are known plant pathogens affecting cereals and leafy greens (32, 33), no symptoms of plant disease were observed in our study, suggesting that the strains present were nonpathogenic or occurred at concentrations too low to cause disease. However, some *Acidovorax* species are beneficial to plants and have strong antibacterial and antifungal activities, aid in nitrification, production of phytohormones, and sensing and transport of organic acids (34, 35, 36).

**FIG 6.**
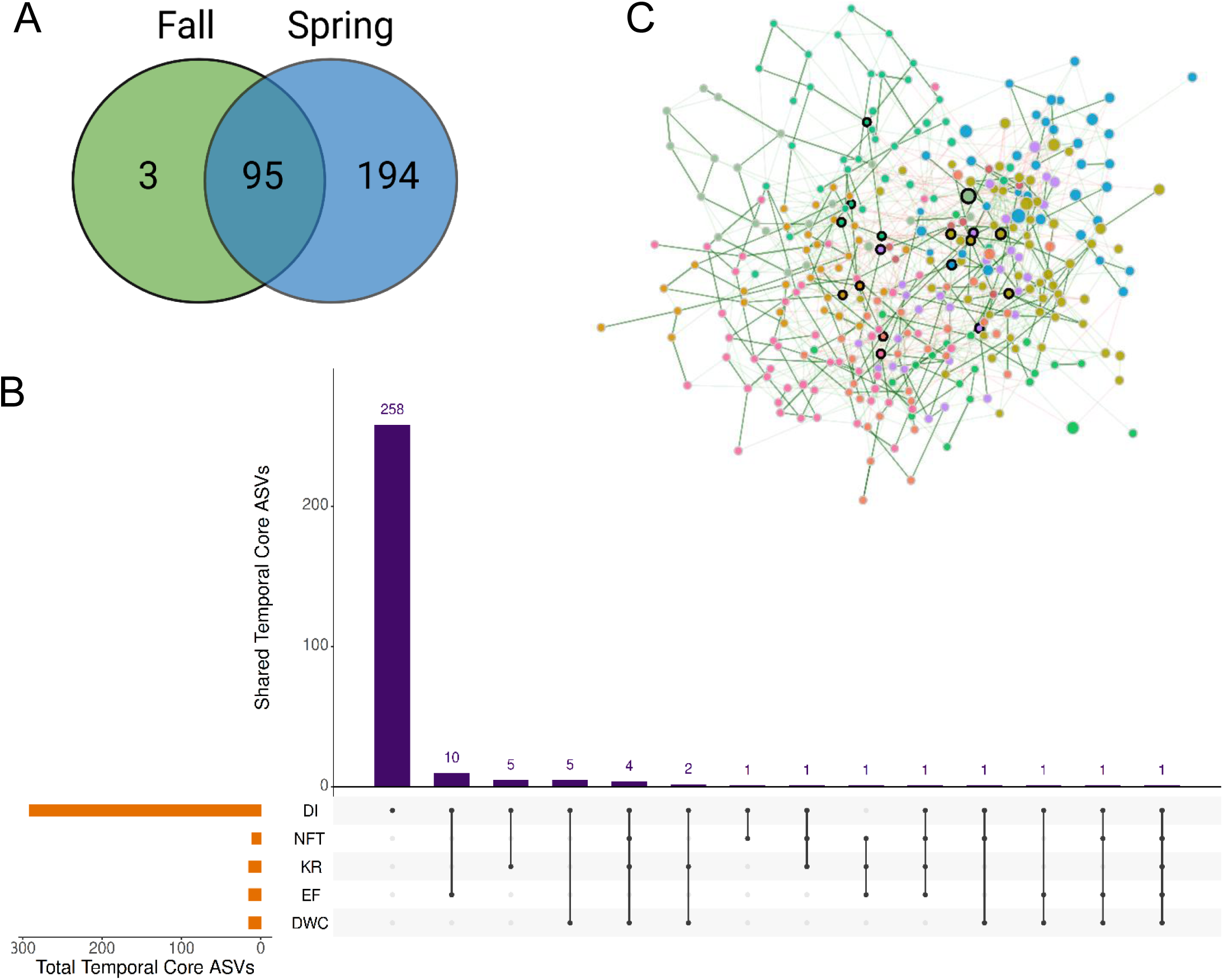
Temporal core ASVs were identified for each system and season as those present in all samples (i.e., occupancy = 1) collected from one system during one season. The resulting system- and season-specific core ASVs were aggregated into a combined set for downstream analyses. Venn diagram shows the number of temporal core bacterial ASVs shared between the two seasons, green for fall and blue for spring (A). UpSet plot shows the number of shared and unique Temporal Core ASVs among the five systems. Each vertical bar represents the number of ASVs present in the intersection of the systems indicated by the connected black dots below. Horizontal bars indicate the total number of ASVs detected in each individual system (B). Networks show bacterial taxa with an occupancy above 0.5 in any system across both seasons. Nodes represent ASVs and are color-coded by network cluster, which is a group of taxa that are more strongly connected to each other than to the rest of the network, as determined by the fast greedy algorithm. These clusters represent putative microbial sub-communities that may share similar ecological niches, respond similarly to system conditions, or participate in related metabolic processes. Nodes marked with a black border line are network hubs, determined as those with the highest betweenness centrality, meaning they serve as key connectors or bridges linking different parts of the microbial network. A higher transparency in the edge color indicates lower association value (C). DI, drip irrigation; DWC, deep water culture, EF, ebb and flow, KR, Kratky, NFT, nutrient film technique.

When the occupancy threshold was relaxed to presence in at least 50% across all samples, 20 high-occupancy ASVs emerged as common members of the nutrient solution microbiota. These included both classified taxa, such as *Devosia, Bdellovibrio, Legionella, Rhodanobacter, Sphingobium, Sediminibacterium, Massilia,* and *Caulobacter*, and unclassified taxa affiliated with the *Sphingobacteriales* and *Sphingomonadaceae. Legionella* was represented by multiple ASVs, underscoring the importance of monitoring taxa with potential relevance to water quality.

Many of the high-occupancy genera identified have also been reported in other hydroponic, aquatic, or plant-associated environments. A study looking at water inputs for hydroponic greenhouses in France found unclassified *Sphingomonadaceae* had the highest taxa abundance in rainwater while *Devosia* and *Bdellovibrio* were more abundant in groundwater sources (37). *Sediminibacterium* had a relative abundance greater than 1% in more than half of the samples collected from an NFT system growing Brassica leafy vegetables in Singapore (18). *Massilia* and *Rhodanobacter* dominated the rhizosphere of hydroponically grown melons in Taiwan and aquaponic systems in South Africa (38, 39). *Caulobacter* was identified in samples from hydroponically grown plants in China (21). In slow sand filtration systems used for greenhouse water treatment, *Legionella* species have constituted a substantial portion of the resident bacterial community (40).

These high-occupancy genera exhibit several traits which may help explain their persistence across soilless farming systems. Many of these genera are motile, a trait that likely promotes their widespread distribution in aqueous based systems (11, 12, 13, 41). Several genera identified here are also associated with aquatic or rhizosphere environments and may contribute to functions such as nutrient cycling or microbial regulation (42, 43, 44). For instance, *Bdellovibrio* is a known bacterial predator that can influence community structure by targeting Gram-negative bacteria (41), while *Devosia* includes nitrogen-fixing and plant-associated species even in the absence of a traditional soil rhizosphere. *Rhodanobacter* has been identified as a bacterial marker of amino acid regulation in hydroponically grown chives and contributes to nitrification, denitrification, and detoxification of plant-derived toxins (46). *Sphingobium* spp. have been implicated in the degradation of polycyclic aromatic hydrocarbons in hydroponically cultivated sudangrass (24). *Caulobacter* may act as a pathogen antagonist in soilless systems, inhibiting wildfire disease, caused by *Pseudomonas syringae* pv. *tabaci*, and other plant pathogens (21, 47). *Devosia* species possess permeases that enhance environmental sensing and chemotaxis (11) and have been isolated as root and hyphal colonizing bacteria in mycorrhiza-rich substrates (48). The persistence of unclassified *Sphingobacteriales* and *Sphingomonadaceae* emphasizes the metabolic versatility and biofilm-forming potential of soilless farming microbiomes (49, 50).

Previous work in soil-based systems has shown that drip irrigation delivery methods, like those found in our DI system, can substantially influence microbial composition (51). Wang et al., (2022) reported that differences in drip irrigation depths and patterns influenced root-soil-microbe interactions, suggesting that fluid movement plays a key role in selecting for specific microbial assemblages. Although this was a soil-based study, this broader principle supports our finding that differences in nutrient solution flow and distribution across system types can result in distinct bacterial community structures. These observations suggest that fluid dynamics in soilless systems may be an important determinant of bacterial persistence and differentiation. System design and fluid movement shape which bacteria persist in these systems, and also how microbes interact with one another within these communities. Hubs are highly connected taxa that play central roles in microbial interactions within a community. The disruption of key taxa ripples through the community via microbe-microbe interactions and can lead to the collapse of critical ecological functions (52). Unlike abundant taxa, hubs are defined by their extensive connectivity, exhibiting high degree of betweenness centrality (i.e., being the shortest path between other taxa), linking multiple, otherwise weakly connected taxa. This position in a community suggests they may mediate resource exchange, signaling, and cross-feeding, that enhance the community stability and resilience. Hubs often influence nutrient cycling and buffering environmental fluctuations that act as stressors. These taxa are often considered keystone taxa with important ecological roles (53). Our network analysis identified 17 hub ASVs across all samples based on centrality metrics (Fig. 6). These ASVs belonged to the genera *Alicyclobacillus*, *Mycobacterium*, *Acidovorax*, *Dyella*, *Desulfosporosinus*, *Actinoallomurus*, *Chitinophaga*, *Elsterales*, *Nonomuraea*, *Arachidicoccus*, *Peredibacter*, *Legionella*, *Caulobacter*, *Pedobacter*, *Mucilaginibacter*, and *Granulicella*, with *Alicyclobacillus* represented by two distinct ASVs.

Notably, *Acidovorax*, *Legionella*, and *Caulobacter* were identified as both the core and hub taxa, although as different ASVs. This pattern indicates that while these genera are consistently present in the system, different strains may occupy distinct ecological roles. Such ASV-level divergence suggests functional differentiation or ecological specialization within these genera, with some strains acting as highly connected network hubs and others contributing to widespread occupancy.

Functional traits may help explain why many of these taxa serve as hubs. Several genera identified as hubs, including *Alicyclobacillus*, *Mycobacterium*, *Caulobacter*, and *Dyella*, are known to be motile and capable of forming biofilms (54, 55, 56, 57). These hubs are also capable of surface colonization and possess metabolic and ecological traits that can support or regulate other microorganisms within the community. For example, through enzymatic conversion of phenolic compounds, *Alicyclobacillus* participates in aromatic carbon degradation, generating metabolites that may serve as substrates for other microorganisms (58). Members of the genus *Acidovorax* exhibit metabolic versatility, including the ability to reduce metal oxides and perform denitrification through both extracellular and intracellular electron transfer pathways (59). *Caulobacter* also exhibits morphological plasticity, forming filamentous cells under environmental stress that can extend above the biofilm surface and enhance nutrient access, dispersal, and resistance to predation (47). These adaptive behaviors can restructure the physical architecture of bacterial communities and contribute to the persistence and functional centrality of *Caulobacter* in soilless farming environments. These traits promote persistence, surface colonization, and interaction with other microbes, which likely increases their centrality in network structure.

#Figure 7 shows the relative abundance of ASVs averaged across each season for all soilless systems. It includes only taxa present in at least half of the samples (occupancy ≥ 0.5) and with a mean relative abundance greater than 1%. When systems were averaged by season, *Methylophilus* exhibited the highest mean relative abundance across every system in the spring and remained dominant in the DI and EF systems during the fall (Table 1). In contrast, *Eoetvoesia* was the most abundant taxon in the DWC and NFT systems in the fall, while *Acidovorax* was most abundant in the KR system.

**FIG 7.**
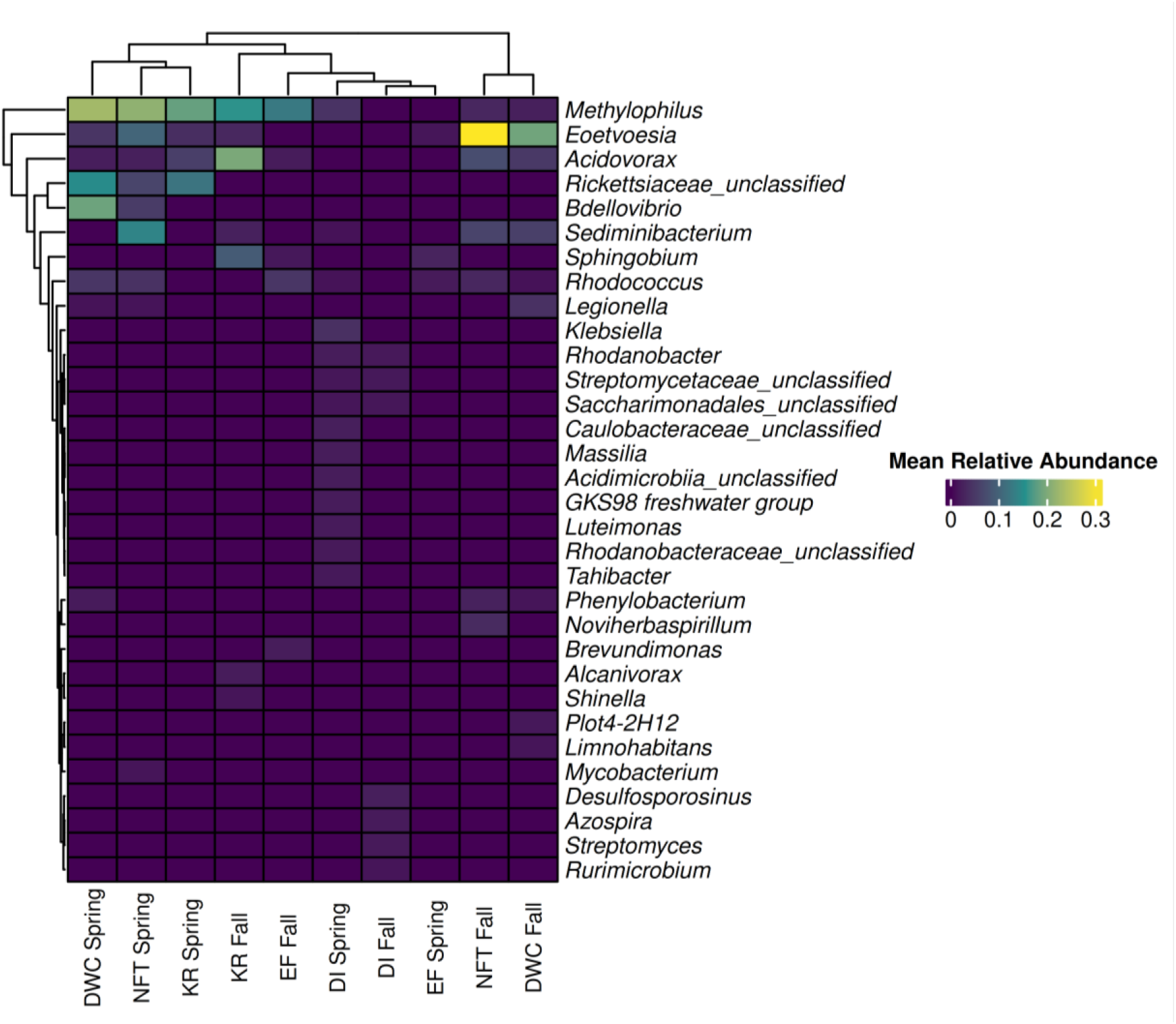
Relative abundance of ASVs averaged over the season for each system. ASVs present in at least half of the samples (i.e., occupancy = 0.8) with an abundance greater than 1% are included. (DI = drip irrigation; DWC = deep water culture, EF = ebb and flow, KR = Kratky, NFT = nutrient film technique).

**TABLE 1.** Genera with highest relative abundances in each system for both seasons. ASVs with occupancy above 0.5 and abundance above 0.005 were included. DI, drip irrigation; DWC, deep water culture, EF, ebb and flow, KR, Kratky, NFT, nutrient film technique.

| Soilless system | Season | Genus | Mean relative abundance (%) | Mean occupancy (%) |
| --- | --- | --- | --- | --- |
| DI | Fall | *Methylophilus* | 12.9 ± 0.12 | 75 ± 0.43 |
|  | Spring | *Methylophilus* | 4.3 ± 0.30 | 100 ± 0.00 |
| DWC | Fall | *Eoetvoesia* | 19.2 ± 0.12 | 100 ± 0.00 |
|  | Spring | *Methylophilus* | 22.9 ± 0.23 | 100 ± 0.00 |
| EF | Fall | *Methylophilus* | 12.6 ± 0.12 | 85.7 ± 0.35 |
|  | Spring | *Methylophilus* | 18.5 ± 0.23 | 66.7 ± 0.47 |
| KR | Fall | *Acidovorax* | 19.8 ± 0.20 | 100 ± 0.00 |
|  | Spring | *Methylophilus* | 18.4 ± 0.07 | 100 ± 0.00 |
| NFT | Fall | *Eoetvoesia* | 30.9 ± 0.24 | 85.7 ± 0.35 |
|  | Spring | *Methylophilus* | 21.4 ± 0.13 | 100 ± 0.00 |

Integrating relative abundance patterns with occupancy data provided insight into how microbial composition shifts across systems and seasons (Fig. S2-S11). *Methylophilus* consistently dominated communities in the spring and remained abundant in DI and EF during the fall, whereas *Eoetvoesia* was more prominent in the fall communities of DWC and NFT. *Acidovorax* was the prevailing taxon in the KR system during fall. Notably, although *Acidovorax* appeared in both the core and hub taxa lists, *Methylophilus* and *Eoetvoesia* did not, despite their high relative abundance. Although *Methylophilus* and *Eoetvoesia* reached high relative abundance, their absence from core and hub taxa suggests that these genera may respond opportunistically to shifts in root-derived carbon inputs rather than occupying persistent ecological roles. Root exudation is known to vary with plant growth conditions and environmental context, and changes in exudate composition can rapidly restructure rhizosphere communities (60). Neighbor-induced exudate shifts in soil systems demonstrate how sensitive microbial communities are to plant metabolic signals, and similar fluctuations in soilless systems may explain why some taxa bloom transiently but do not achieve widespread occupancy or network centrality (19, 61).

## Methods

### Experimental location and soilless systems setup

The study was conducted at the Penn State Greenhouse Facility at University Park, PA, comparing side-by-side under controlled environmental conditions five alternative soilless growing systems, including DWC, KR, NFT, DI, and EF soilless growing systems, using bok choy (*Brassica rapa* subsp. *chinensis*) cv ‘Li Ren Choi’ (Johnny’s Selected Seeds, Winslow, ME, USA) as a test crop. Each soilless growing system was built as a modular system with a footprint of 1.44 m^2^ (1.2 × 1.2 m), and 36 plants per unit were planted, establishing a density of 25 plants/m^2^ across all soilless systems. Soilless systems were arranged in a randomized complete block design with three replications. The study was repeated twice, in the fall-winter (November - December) of 2023 and the winter-spring (February – March) of 2024. The alternative soilless systems tested were selected for their popularity and adoption at commercial scale and for their unique design features. As described by Blunk et al. (2023) and Poudel and Di Gioia (2025), the DWC systems were set up using large (135×135×17 cm) black trays (Botanicare, Vancouver, WA) sustained by metallic frames. The trays served as reservoir tanks and were filled with nutrient solution. Styrofoam (expanded polystyrene) floating boards (rafts) were cut to fit each tray. Planting holes, each with a small net pot used to hold in place the bok choy seedlings, were set into each raft in six staggered rows (20 cm apart) to establish the defined plant density (25 plants/m^2^). An air pump (60 L/min, Vivosun, Ontario, CA, USA) with four air stones per module was used to continuously aerate the nutrient solution. The KR system differed from the DWC system, only for two aspects: i) the styrofoam board did not float on the nutrient solution but rather was suspended at the top of the tray filled with nutrient solution, and ii) the nutrient solution was not aerated with the air pump, instead, as the nutrient solution was consumed it was not replenished and the plants were oxygenated through the air gap created between the surface of the nutrient solution and the styrofoam rafts holding the plants. Both the DWC and KR systems can be classified as static hydroponic systems. The modular NFT systems (CropKing Inc., OH, USA) had the same footprint as the other soilless growing systems and were built with six 1.2-m-long food-grade PVC channels, mounted on a metallic frame with a slight slope (4%). Each growing channel had six (2.5 × 2.5 cm) holes 20 cm apart, in which bok choy seedlings were positioned. The nutrient solution serving the NFT systems was kept underneath each module in a 94 l black reservoir tank containing a submersible water pump (Active Aqua, AAPW250, Fairfield, CA, USA) that continually served each growing channel. The drained nutrient solution was recollected in the same reservoir tank by gravity, and the nutrient solution was oxygenated while falling from the PVC collector pipe placed at the lower end of the NFT growing channels, into the reservoir tank. The NFT system can be classified as a recirculating hydroponic growing system.

The DI and EF systems were set up using the same large trays and metallic frames used for the DWC systems; however, in this case, bok choy plants were grown in pots (black) of 1.45 L (Koba corporation, Middlesex, NJ, USA) with holes at the bottom filled with a peat-perlite growing media mix (PRO-MIX BX™, Premier Tech Ltd., Rivière-du-Loup, Canada) and placed in six rows of six pots establishing the same plant density of the other systems. The EF systems had a 220 l reservoir tank placed underneath each growing tray, and the trays were flooded with nutrient solution using a submersible water pump activated through a timer, for 5 minutes, once a day. The nutrient solution was absorbed by the pots through subirrigation, and the excess was drained and recollected in the same reservoir tank underneath each EF growing module.

In the case of the DI system, the setup was very similar to the EF system; however, the nutrient solution was delivered by auto-compensating drippers with a flow rate of 1 l/h. A single dripper served two pots, and fertigation events were scheduled with a timer twice a day. The excess of nutrient solution was drained and collected into a bucket placed at the bottom of each module but was not recirculated. The EF and DI can both be classified as substrate-based soilless growing systems but have different nutrient solution delivery methods and management. In the EF growing systems, the nutrient solution was delivered from the bottom of the pots and absorbed by the growing media by capillarity with an upward movement, and the drained nutrient solution was recirculated according to a closed-cycle management approach. Instead, in DI systems, the nutrient solution was delivered at the top of the growing media and moved downward by gravity, and the drained nutrient solution was not recirculated, but was managed according to an open-cycle management scheme.

### Crop and growing system management

Bok choy was seeded in rockwool cubes (2.5×2.5×4-cm, Grodan, Roermond, The Netherlands) on October 24, 2023, and January 23, 2024, for the fall and spring cycles, respectively, and was transplanted into each modular growing system 17-18 days after seeding, on November 9, 2023, and February 9, 2024, respectively.

As described by Poudel and Di Gioia (2025), bok choy plants were nourished with a standard modified Sonneveld nutrient solution containing macro- and micro-minerals at the following concentrations: 153 N, 31 P, 210 K, 100 Ca, 24 Mg, 59.6 S, 1 Fe, 0.25 Mn, 0.14 Zn, 0.03 Cu, 0.16 B, and 0.04 Mo mg/L. During the seedling production phase, a half-strength nutrient solution was applied, while starting at planting, the same nutrient solution was applied to all the soilless growing systems. The nutrient solution was prepared using reverse-osmosis water and greenhouse-grade water-soluble fertilizers such as calcium nitrate, ammonium nitrate, potassium nitrate, potassium sulfate, magnesium sulfate, monopotassium phosphate, Fe-EDTA, Mn-EDTA, boric acid, copper sulfate, and sodium molybdate. Two concentrated (100 times) stock nutrient solutions (A and B) were prepared in 50 L tanks to keep separated Ca from S and P fertilizers and were mixed and delivered at the final dilution to each soilless growing system through volumetric nutrient solution injectors (Dosatron^®^, Clearwater, FL, USA). The pH and electrical conductivity (EC) of the nutrient solution were checked upon mixing the nutrient solution and were monitored weekly using a portable pH-EC meter (HI991300, Hanna Instruments Inc., Woonsocket, RI, USA) to keep pH (5.8-6.2) and EC (1.8-2.2 dS/m) within the optimal range, in both experiments. The same pH-EC meter was also used to measure the nutrient solution temperature. The nutrient solution of each modular soilless setup was recirculated in a closed system, except for the DI modules that were managed as open-cycle systems. Adjustments of pH and EC were made by topping up the recirculating nutrient solution to replenish the water consumed, using fresh nutrient solution, and when needed, pH adjustments were made using small quantities of pH-Up^©^ (General Hydroponics, USA), a solution containing potassium hydroxide and potassium carbonate. For the water culture systems (DWC, KR, and NFT), dissolved oxygen (DO) concentrations were monitored weekly using an A223 RDO/DO meter (Orion Star Inc., Waltham, MA, USA).

Bok choy plants were harvested on December 15, 2023, and March 15, 2024, at 36 and 35 days after planting (DAP) in the fall-winter and winter-spring cycle, respectively. During the two crop cycles, the average air temperature in the greenhouse was maintained around 22-23 °C. Relative humidity ranged between 30% and 85% between the two growing seasons. An integrated crop management approach with minimum agrochemical applications was used to manage key pests such as thrips (*Frankliniella occidentalis*) and aphids (*Myzus persicae*) during both crop cycles.

### Nutrient Solution and Bok Choy Sampling

After the bok choy was transplanted, one nutrient solution sample was taken from each of the 15 modular soilless systems weekly. The nutrient solution from the DWC and KR systems was collected directly from the reservoir. The NFT nutrient solution was sampled from the drain which flowed back into the reservoir. The nutrient solution from the DI and EF systems was collected from the drain immediately after the system had been fertigated. Sterile bottles, autoclaved and wrapped in aluminum foil, were used to collect nutrient solution samples from each system. At each sampling point the foil was removed and the bottle was dipped into the nutrient solution without allowing contact between the solution and the operator’s gloves. Gloves were disinfected with ethanol between samples to minimize chances of cross-contamination. After all samples were collected, they were transported from the greenhouse to the lab for analysis within one hour of collection. At the end of the two growth cycles, one bok choy plant was randomly selected from each system and harvested using disinfected shears. Twenty-five grams of outer leaves were removed and placed into a sterile Whirl-Pak bag (Whirl-Pak, Pleasant Prairie, WI).

### Enumeration of Aerobic Bacteria

The bok choy leaves were placed in a sterile Whirl-Pak bag (Whirl-Pak) with 225 mL of sterile 1 × phosphate-buffered saline (PBS) and homogenized in a stomacher for 1 minute at 230 RPM. The nutrient solution and homogenate from the leaves were vortexed, serially diluted, and plated in duplicate onto aerobic plate count (APC) Petrifilms (3M, Hutchinson, MN) in the fall. In the spring, standard plate count agar (SPCA) was used for aerobic bacteria due to a shortage of APC Petrifilms. APC Petrifilms and SPCA plates were incubated at 35°C for 48 ± 2 h and enumerated to determine the concentration of aerobic mesophilic bacteria per milliliter of nutrient solution or gram of bok choy.

### DNA Extraction

For each soilless system, 100 mL of nutrient solution from each of the biological replicate samples wercombined to create a composite sample representative of a given system at a given time point. The composite samples were filtered through 0.45µm Nalgene Rapid-Flow Sterile Disposable Filters (Thermo Fisher Scientific, Waltham, MA, USA). DNA was extracted directly from the filters using the DNeasy PowerWater Kit (Qiagen, Hilden, Germany) according to the manufacturer’s instructions.

To assess potential contamination introduced during DNA extraction, negative extraction controls consisting of nuclease-free water processed through the extraction kit were included. In addition, the ZymoBIOMICS Microbial Community Standard (Zymo Research, Irvine, CA, USA) was processed in parallel as a positive control to verify extraction efficiency and downstream sequencing performance.

### 16S rRNA V4 Amplicon Sequencing

Extracted DNA was used to prepare libraries in the One Health Microbiome Center Research Collaboratory using their standard protocols (62). Briefly, the 16S rRNA V4 region was amplified using the KAPA HiFi HotStart mastermix and 515F (GTGYCAGCMGCCGCGGTAA) and 806R (GGACTACNVGGGTWTCTAAT) primers with partial overhangs for the i7 and i5 adapters (63). Primary PCRs were conducted in 10 µL reactions with KAPA HiFi hot start enzyme and SYBR green to monitor reaction progress. Samples were subsequently diluted 100× before indexing with unique dual 10nt indexes based on the Illumina Tagmentation Sets A–D. Resulting indexed libraries were quantified with PicoGreen and pooled at equimolar concentrations. The library was size-selected by gel purification followed by Ampure bead cleanup using a 0.6× ratio. The DNA was sequenced on an Illumina NextSeq 2000 with XLEAP P1 600-cycle reagents for both in vitro and in vivo samples, generating paired-end reads of approximately 270 bp after demultiplexing. Primer sequences were removed allowing for an error rate of 0.15 using Cutadapt/ QIIME2 version 2023.5, discarding any read without a valid primer sequence on both ends.

### Sequence Data Analyses

Demultiplexed Illumina paired-end reads were processed using the DADA2 pipeline (v1.26.0) in R. Reads were filtered and trimmed using the filterAndTrim() function with the following parameters: truncLen = c(240, 240), trimLeft = c(0, 0), maxN = 0, maxEE = c(2, 2) (expected error threshold), truncQ = 3, rm.phix = TRUE, compress = TRUE, and multithread = TRUE. Error rates were learned from the filtered reads using learnErrors() with default settings. ASVs were inferred using the dada() function (pool = FALSE), and paired-end reads were merged using mergePairs() with default parameters. Following merging, only sequences between 251–256 bp were retained, corresponding to the expected length of the V4 region of the 16S rRNA gene. Chimeras were removed using removeBimeraDenovo (method = “consensus”, multithread = TRUE).

A sequence table was constructed using makeSequenceTable(). Taxonomy was assigned with assignTaxonomy() using the SILVA v138.2 reference database, followed by species-level refinement with addSpecies(). ASVs annotated as chloroplast or mitochondria were removed before downstream analyses.

### Statistical Analysis

All statistical analyses were completed in R version 4.4.1. A linear model was used to assess the effects of system, temperature, and pH on log-transformed aerobic mesophilic bacteria concentrations. Separate models were run for each season, and interaction terms were included to evaluate whether the relationships between environmental variables and APC differed by season. Temperature and pH were treated as continuous covariates, and system was treated as a categorical fixed effect. When system effects were significant, post hoc pairwise comparisons were conducted using Tukey’s Honestly Significant Difference HSD to identify differences between individual systems. Aitchison distances were computed from CLR-transformed ASV data. Because CLR transformation cannot be applied to zeros, raw ASV counts were first processed with the count zero multiplicative (CZM) method using cmultRepl (zCompositions; label = 0, method = “CZM”, output = “p-counts”, z.warning = 0.99), and the resulting values were converted to relative abundances by closing each sample to sum to 1. ASVs with extremely low abundance were removed by retaining only those taxa with a minimum relative abundance > 1×10⁻⁶ across all samples prior to CLR transformation. Pairwise community differences based on Aitchison distance were then evaluated using the pairwiseAdonis2 function (pairwiseAdonis package v. 0.4.1). Differentially abundant ASVs between system groups were identified using the aldex.clr and aldex.ttest functions from the ALDEx2 v1.30.0 package. ASV reads were rarified to a depth of 1,500 reads prior to imputation or compositional transformations, and Shannon Diversity was computed and used to assess diversity over time, between systems, and within systems.

Core taxa analysis code was adapted from Rolon et al (2023). A phyloseq object was constructed from the ASV table, taxonomy table, and sample metadata, and then subset by system (DI, DWC, EF, KR, NFT) and season (spring, fall). Within each system–season subset, ASVs with a total count of zero across all samples were removed. For occupancy, ASV tables were converted to presence–absence, and occupancy was defined as the proportion of samples in which each ASV was detected. To obtain mean relative abundances while accounting for zeros, count data were first processed with the count zero multiplicative (CZM) replacement method (cmultRepl, method = “CZM”, z.warning = 0.99, output = “p-counts”; zCompositions package), and then transformed to compositions closed to 1 using Aitchison geometry (acomp; compositions package). Mean relative abundance for each ASV in each system–season combination was calculated as the average relative abundance across all samples in that subset. For sample-level analyses, compositional ASV tables were similarly generated for each system–season combination and then combined into a single dataset annotated with system, season, and sampling day. Alpha diversity was computed from data pre-CZM treatment using the vegan package, with observed richness calculated via specnumber() and Shannon and Simpson indices via diversity(index = “shannon”) and diversity(index = “simpson”), respectively.

All code used for sequence processing, statistical analyses, and figure generation is publicly available in a dedicated GitHub repository (https://github.com/abywats717/Bok_Choy_Soilless_Farming). Raw demultiplexed sequence data have been deposited in the NCBI Sequence Read Archive under BioProject accession number PRJNA1366713.

## Acknowledgments

This research was supported by the Pennsylvania Department of Agriculture grant C940001528, the USDA NIFA and Hatch Appropriation under Project PEN04853 and Accession 7005519, and the Multistate project 4666, as well as Lester Earl and Veronica Casida Career Development Endowment. FD’s contribution was supported by the USDA National Institute of Food and Agriculture and Hatch Appropriations under Project #PEN05002, Accession #7007517. Amplicon libraries were prepared by the Penn State One Health Microbiome Center Research Collaboratory under the guidance of Dr. Jordan Bisanz. Amelia Navarre, Erin Readinger, and Kailee Shotto assisted with media preparation, sample collection, and processing. Dr. Laura Rolon assisted with microbiota analysis.

